# Assessing vaccine-mediated protection in an ultra-low dose *Mycobacterium tuberculosis* murine model

**DOI:** 10.1101/2023.03.22.533820

**Authors:** C.R. Plumlee, H.W. Barrett, D.E. Shao, K.A. Lien, L.M. Cross, S.B. Cohen, P.T Edlefsen, K.B. Urdahl

**Affiliations:** Center for Global Infectious Disease Research, Seattle Children’s Research Institute, Seattle, WA, 98109, USA; University of Washington, Dept. of Global Health, Seattle, WA, 98109, USA; Vaccine and Infectious Disease Division, Fred Hutch Cancer Center, Seattle, WA, 98109, USA; University of Washington, Dept. of Immunology, Seattle, WA, 98109, USA; University of Washington, Dept. of Pediatrics, Seattle, WA, 98109, USA

**Author notes:** Equal contributions (C.R. Plumlee, H.W. Barrett).

## Abstract

Despite widespread immunization with Bacille-Calmette-Guerin (BCG), the only currently licensed tuberculosis (TB) vaccine, TB remains a leading cause of mortality globally. There are many TB vaccine candidates in the developmental pipeline, but the lack of a robust animal model to assess vaccine efficacy has hindered our ability to prioritize candidates for human clinical trials. Here we use a murine ultra-low dose (ULD) *Mycobacterium tuberculosis* (Mtb) challenge model to assess protection conferred by BCG vaccination. <u>We show that BCG confers a reduction in lung bacterial burdens that is more durable than that observed after conventional dose challenge, curbs Mtb dissemination to the contralateral lung, and, in a small percentage of mice, prevents detectable infection.</u> These findings are consistent with the ability of human BCG vaccination to mediate protection, particularly against disseminated disease, in specific human populations and clinical settings. Overall, our findings demonstrate that the ultra-low dose Mtb infection model can measure distinct parameters of immune protection that cannot be assessed in conventional dose murine infection models and could provide an improved platform for TB vaccine testing.

## INTRODUCTION

New and effective tuberculosis (TB) vaccines are urgently needed. Although BCG vaccination can provide protection in infants and older children in some clinical settings^1–4^, it has proven to be inadequate to combat the global pandemic. There are numerous TB vaccine candidates in the developmental pipeline^5^, but it will not be feasible to conduct human efficacy trials for most of them. The standard for assessing vaccine efficacy is prevention of disease, which occurs in only a small percentage of infected individuals, resulting in dauting sample sizes and costs needed to complete human trials. Unfortunately, identifying promising vaccine candidates to prioritize for human efficacy trials has been hindered by the lack of reliable animal models that adequately predict human TB vaccine efficacy^6, 7^.

Historically, mice have been the most commonly used model for testing TB vaccine candidates due to their ease of use, cost-effectiveness, and the relative conservation of the mammalian immune system. However, limitations in the current mouse model, in which mice are infected with ∼50-100 CFU by aerosol, have shaken confidence regarding how well findings in mice can be translated to humans^6, 7^. There is minimal variability in the performance of different vaccines in the current model. Most TB vaccines confer ∼1 log of protection, reducing the lung burden from ∼10^6^ to ∼10^5^ CFU, but only if assessed 4-6 weeks after aerosol challenge. This protection is transient and usually dissipates by 3-4 months post-infection (p.i.)^8, 9^. Furthermore, it is unclear if the vaccine-induced mechanisms that enable mice to transiently reduce their bacterial burdens in the setting of an ultimately unsuccessful immune response are relevant to the types of immunity required for long lasting protection against the clinical manifestations of human TB. Because mice in this model are unable to eradicate, or even durably control Mtb, some have suggested that mice may lack the fundamental immune effector molecules needed for Mtb control^10^. The failure of mouse vaccine testing to reliably predict results in human TB vaccine trials have reinforced these concerns about the relevance of the mouse model for TB vaccine testing^6, 7^.

Recently we developed a physiologic ultra-low dose (ULD) (i.e., 1-3 CFUs) Mtb mouse infection model that more closely resembles several aspects of human Mtb infection^11^. Here we test the ability of the ULD challenge model to assess immunity conferred by BCG vaccination, the vaccine for which the most human efficacy data exists. In contrast to the conventional dose model, we show that BCG-vaccinated mice challenged with 1-3 Mtb CFU exhibit a more durable reduction in lung bacterial burdens. Vaccinated mice had an increased ability to contain infection to a single lung and prevent Mtb dissemination to the contralateral lung. Finally, vaccinated mice exhibited a significantly higher proportion of animals with no detectable infection. Thus, the ULD model provides a promising platform for TB vaccine testing, as it affords the opportunity to measure distinct parameters of vaccine-mediated immunity that cannot be assessed in the current mouse model.

## RESULTS

### BCG-mediated reduction of lung bacterial burdens is transient in the current mouse model

The capacity of BCG to mediate protection in C57BL/6 (B6) mice infected with 50-100 Mtb CFU has been assessed by many labs, but we sought to repeat this experiment in our own hands to directly compare the conventional dose with the ULD model. We subcutaneously immunized B6 mice with 10^6^ BCG-Pasteur 8 weeks prior to aerosol challenge with H37Rv Mtb and determined the lung bacterial burdens at days 42 and 120 post-infection. As previously shown^8, 9^, BCG immunization provided about one log of protection against lung bacterial burden at day 42, but this protection was transient and dissipated at later timepoints; there was no significant difference in bacterial burdens between unimmunized and BCG-immunized mice at day 120 post-infection (**Figure 1)**.

**Figure 1.**
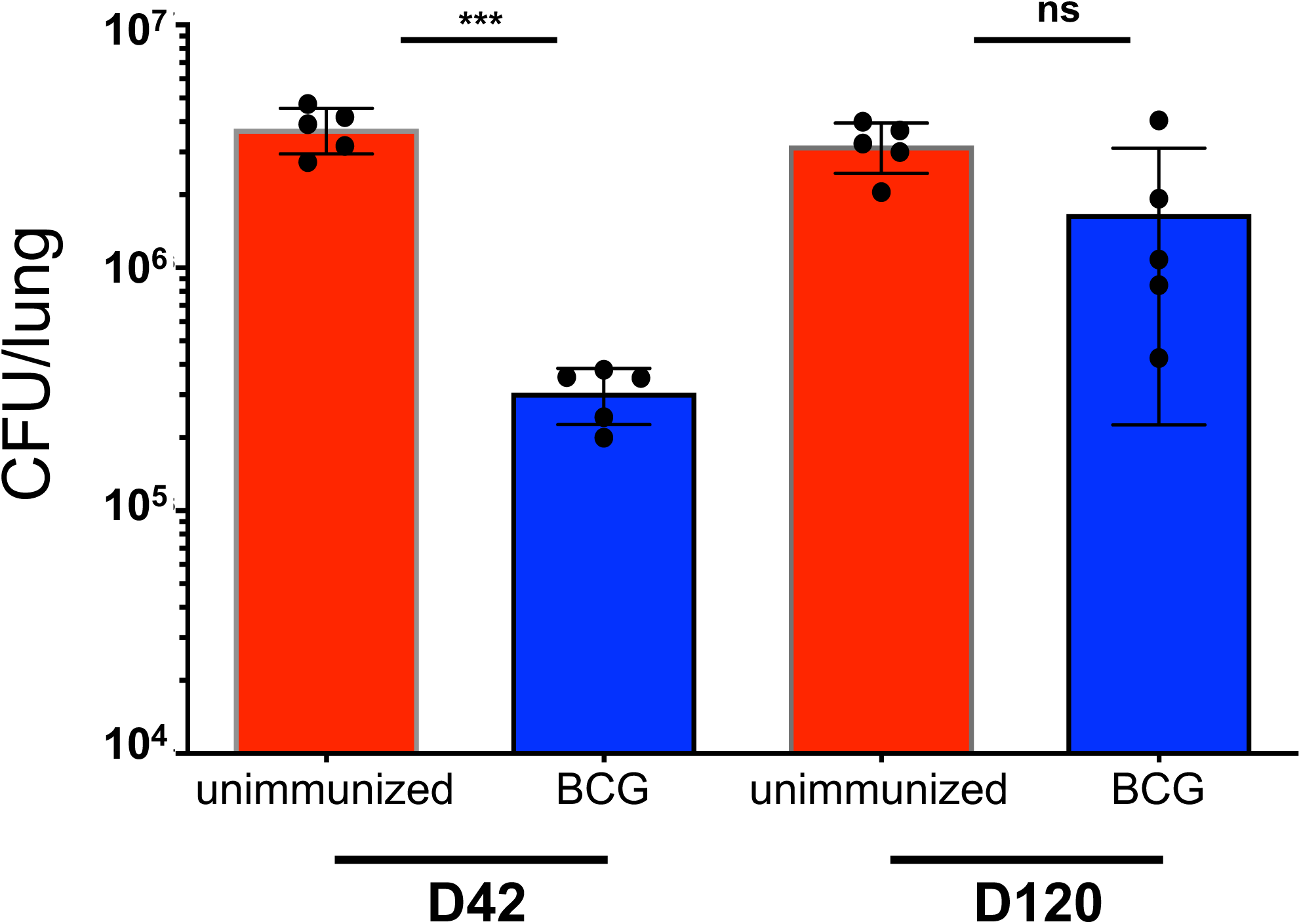
BCG-mediated reductions in Mtb lung burdens are not durable in a conventional-dose infection. C57BL/6 (B6) mice were aerosol infected with a conventional dose (CD) (50-100 CFU) of H37Rv Mtb eight weeks following either subcutaneous (s.c.) immunization with 10^6^ BCG-Pasteur (BCG) or no immunization (unimmunized). On day 42 or 120 post-infection, CFU were enumerated from lung homogenates plated onto 7H10 plates. These data represent 5 mice per group and are shown as mean ± SEM. Single-group comparisons were analyzed using an unpaired t test. ***p<0.001.

### Measuring vaccine-mediated immunity in the ULD Mtb model

Next, we assessed the efficacy of BCG in the ULD aerosol Mtb challenge model, a model in which the aerosolized dose is reduced with a goal of infecting only ∼60-80% of the mice in the infection chamber^11^. As previously described using an ULD infection of a pool of bar-coded Mtb strains, the number founding strains detected using bar-codes after ULD infection approximates a Poisson distribution; most mice are infected with a single founding Mtb strain, whereas fewer are infected with two or three founding strains^11^. **Figure 2** depicts a BCG immunization experiment in which we assessed the bacterial burdens in the right and left lungs separately and in the spleen 9 weeks after aerosol ULD challenge. Amongst those mice with detectable infection (>0 CFU), we observed that, compared to unimmunized mice, BCG-immunized mice had lower overall lung bacterial burdens (pooling right and left lungs) (**Figure 2A**), and lower spleen bacterial burdens (**Figure 2B**). We also observed that 7/20 of the unimmunized mice and 10/20 of the BCG-immunized mice had no detectable infection in either lung (**Figure 2A**), a difference that was not statistically significant in this single experiment (p=0.53). While the same seven mice with undetectable lung bacterial burdens also had no detectable splenic bacterial burdens, two mice that did have detectable lung infection in the BCG-immunized group failed to show evidence of splenic bacterial burdens (12/20 BCG-immunized mice had no recoverable Mtb from their spleen, compared to 10/20 from the lungs). <u>Thus, BCG immunization may prevent Mtb dissemination to the spleen or promote splenic Mtb clearance.</u>

**Figure 2.**
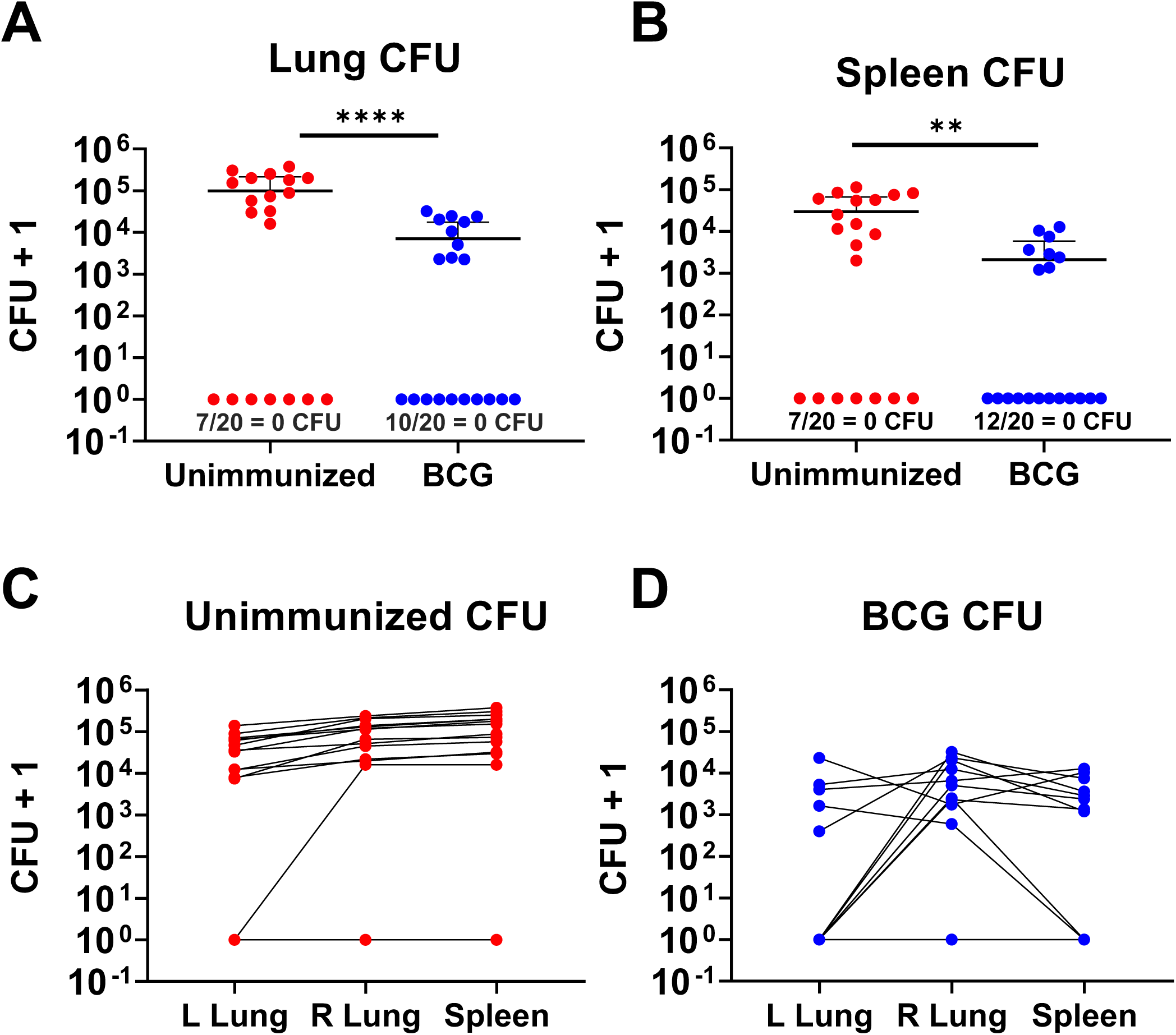
Assessing BCG efficacy in the ultra-low dose Mtb model. B6 mice were aerosol infected with an ULD (1-3 CFU) of H37Rv Mtb 8 weeks following either s.c. immunization with 10^6^ BCG-Pasteur (n=20) or no immunization (n=20). On day 63 post-infection, CFU were enumerated from left lung, right lung, or spleen homogenates plated onto 7H10 plates. **A)** Combined lung CFUs or **B)** spleen CFUs from unimmunized and BCG-immunized mice are graphed. Counts from left lungs, right lungs and spleen are graphed separately from, **C)** unimmunized mice or **D)** BCG-immunized mice. There were 20 mice per group, and the data are graphed as mean ± SD. Single-group comparisons were analyzed using an unpaired t test <u>and excluding mice with 0 CFU</u>. **p<0.01, ****p<0.0001.

To further assess the capacity of BCG immunization to restrict Mtb dissemination, we also assessed bacterial burdens separately in the right and left lungs. In the ULD model, since infection is usually established by a single founding Mtb strain, the lungs are often infected unilaterally; most commonly, the right lung is infected because the right lung and bronchus are larger than the left ^11^. Our previous ULD studies using bar-coded Mtb strains showed that bilateral lung infection usually represents the dissemination of a single Mtb strain from the infection-seeded lung to the contralateral lung. In the experiment shown in **Figure 2**, we observed that 5/10 of the BCG-immunized mice exhibited unilateral lung infection compared to only 1/13 of the infected unimmunized mice (p= 0.023; **Figure 2C and D**). Of the two BCG-immunized mice with pulmonary Mtb infection but no detectable splenic bacterial burdens, one was infected in the right lung only while the other had bilateral lung infection (**Figure 2D**). Thus, the ULD model has the capacity to assess a vaccine’s ability to prevent dissemination, a parameter of protection that cannot be assessed in conventional dose infections because dissemination occurs in all mice regardless of vaccination status.

### BCG confers durable reductions in lung bacterial burdens in ULD-challenged mice

As previously shown^8, 9^, and as demonstrated in **Figure 1**, BCG-mediated reductions in lung bacterial burdens are abrogated by 100-120 days post-challenge. To assess the durability of BCG-mediated protection in the ULD model, we performed a time course and assessed bacterial burdens in the lungs and spleen of ULD-infected mice at an early (d14), intermediate (d42), and late timepoint (d115). <u>Because ULD challenge results in some mice with no detectable lung bacteria, even in the absence of immunization, we assessed bacterial burdens only in mice with detectable bacterial burdens and performed a separate analysis to determine the proportion of mice with undetectable bacteria (the latter is shown in</u> **Figure 5**). At day 14 post ULD-infection, bacterial burdens were similar in both unimmunized and BCG-immunized infected mice, but at days 42 and 115, bacterial burdens were reduced by approximately one log (p=0.002 and p<0.001, respectively, <u>excluding those with 0 CFU</u>; **Figure 3A**). These data suggest that BCG-mediated reductions of lung bacterial burdens are more durable in ULD-infected mice compared to conventional dose-infected mice, but the intentional heterogeneous nature of the model makes it difficult to draw conclusions based on a single experiment.

**Figure 3.**
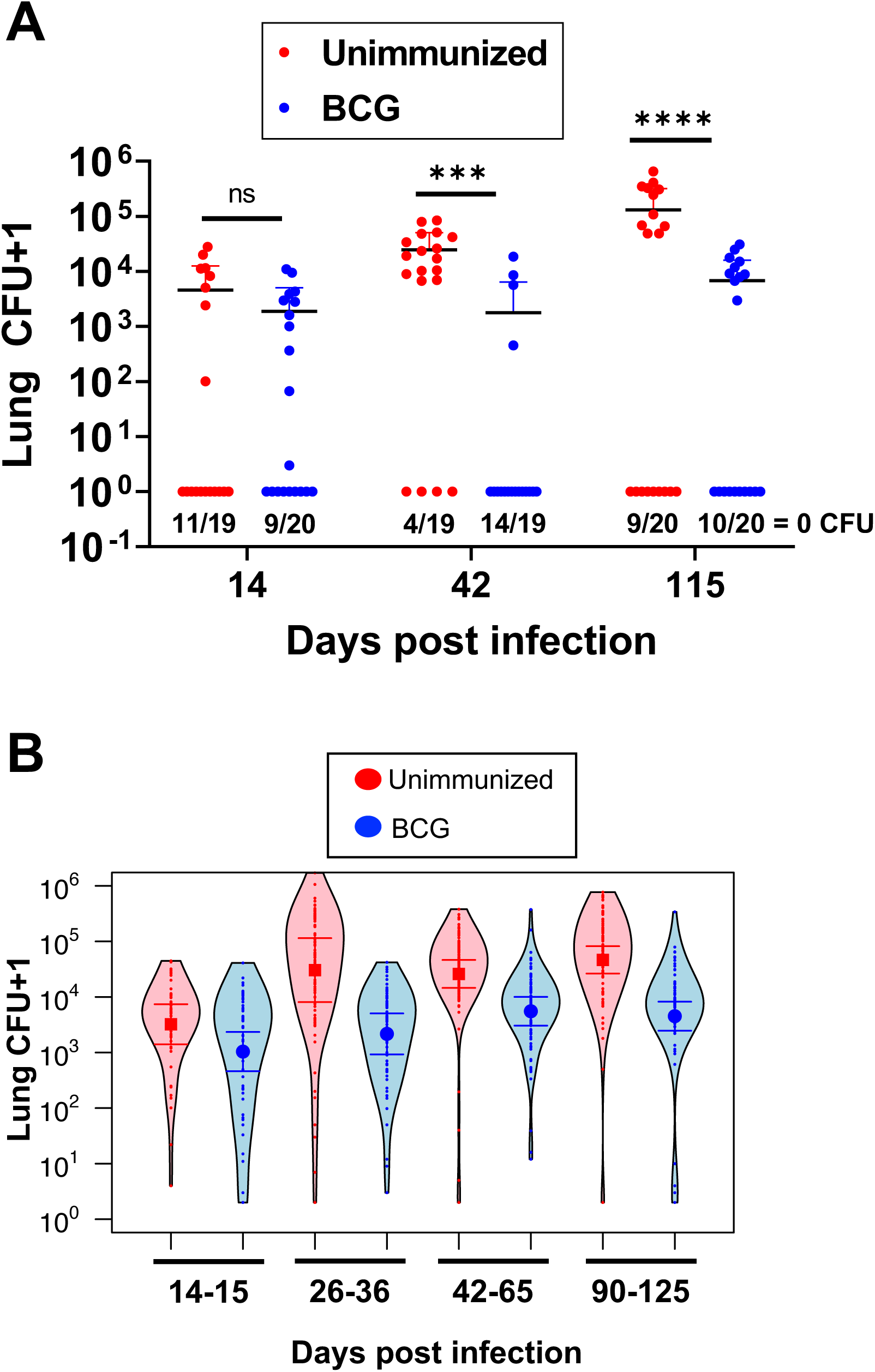
BCG-mediated reductions in Mtb lung burden are <u>more</u> durable in the ULD model. **A)** Combined lung CFU from a single experiment time course of ULD-infected B6 mice with or without BCG immunization. Combined lung CFU were enumerated on days 14, 42, and 115 post-infection. There were 19 or 20 mice per group, and the data are graphed as mean ± SD. Single-group comparisons were analyzed using an unpaired t test (<u>excluding mice with 0 CFU</u>). ***p<0.001, ****p<0.0001. **B)** Combined lung CFU from a compilation of 31 experiments, <u>excluding mice with 0 CFU</u>, separated by timepoint post-infection of ULD-infected B6 mice with or without BCG immunization. Error bars are 95% confidence intervals in a fixed effects negative regression model. All timepoints except days 14-15 post-infection have a p < 0.001.

Next, we performed multiple ULD challenge experiments comparing BCG immunized vs. unimmunized mice to rigorously assess the reproducibility of our findings. **Supplemental Table 1** shows results from 31 individual experiments assessing BCG efficacy in the ULD model, representing a total of 537 unimmunized and 543 BCG-immunized mice with lung bacterial burdens assessed at timepoints ranging from days 14-125 post-infection. **Figure 3B** shows the combined (right and left lungs pooled) bacterial burdens from all experiments binned by similar timepoints post-infection. Similar to prior reports using a conventional dose Mtb challenge^12, 13^, BCG had no effect on bacterial burdens at the earliest timepoints assessed, days 14-15 post-infection. However, BCG-immunized mice had lower lung bacterial burdens than unimmunized mice at every later timepoint (<u>excluding those with 0 CFU</u>). This reduction remained robust (∼1 log) even at days 90-125 post-infection. Thus, unlike the transient reduction of bacterial burdens observed in the conventional dose Mtb challenge model^8, 9^, BCG durably reduces the lung bacterial burdens in ULD-challenged mice for at least 4 months post-infection.

### BCG prevents Mtb dissemination to the contralateral lung

To better assess the reproducibility of BCG’s capacity to prevent dissemination, we assessed the proportion of mice with bilateral vs. unilateral lung infection in all experiments shown in **Supplemental Table I**. Based on raw CFU data, there was no difference in the proportion of mice with bilateral lung infection between unimmunized vs. BCG-immunized groups at days 14-15 post-infection. At this early timepoint, the proportion of mice with bilateral lung infection was low even in unimmunized mice, suggesting that dissemination had not yet occurred in either group. This conclusion is supported by the observation that <20% of ULD-infected mice had detectable splenic Mtb burdens at d14 post-infection (data not shown). At all later timepoints, however, the proportion of mice with bilateral lung infection was higher in unimmunized compared to BCG-immunized mice <u>even out to days 90-125 post-infection</u> (**Figure 4A**). Although bilateral lung infection in the ULD model usually reflects Mtb dissemination from the initially infected lung to the contralateral lung, it can sometimes represent separate aerosolized infections by distinct bacilli in each individual lung. To assess true dissemination more accurately, we ULD-infected mice with a pool of 50 bar-coded H37Rv Mtb strains that we have previously characterized^11^. Amplified genomic DNA extracted from bacterial colonies of each infected mouse lung were sequenced to determine the number of unique founding Mtb strains in each lung (**Figure 4B**). Because the probability of separate infections with the same bar-coded strain is only ∼1 in 78 when using the 50 bar-coded Mtb pool in the ULD model^11^, bilateral lung infection with a single Mtb strain in both lungs likely represents true dissemination (e.g. BCG 36L and 36R). In contrast, when mice have different Mtb strains in each lung (e.g. BCG 28L and 28R), this reflects separate infections of each lung. Thus, if dissemination to the contralateral lung is assessed by bacterial burden alone without assessing Mtb bar-codes, the ability of a vaccine to prevent dissemination will be underestimated due to falsely categorizing separate infections in each lung as dissemination events. In this experiment, BCG was deemed to have 50.2% efficacy (p = 0.002); **Figure 4C**) in preventing dissemination when measured as the proportion of mice with bilateral Mtb infection. However, if dissemination was defined as the proportion of mice that possessed at least one identical bar-coded Mtb strain in both lungs, then BCG exhibited an efficacy of 73.5% (p = 0.001, **Figure 4D**). Compiling data from all experiments in which we performed bar-coded infection and sequencing (n=5) revealed that BCG exhibited 79.7% efficacy in preventing dissemination of Mtb to the contralateral lung (p < 0.001; **Figure 4E**).

**Figure 4.**
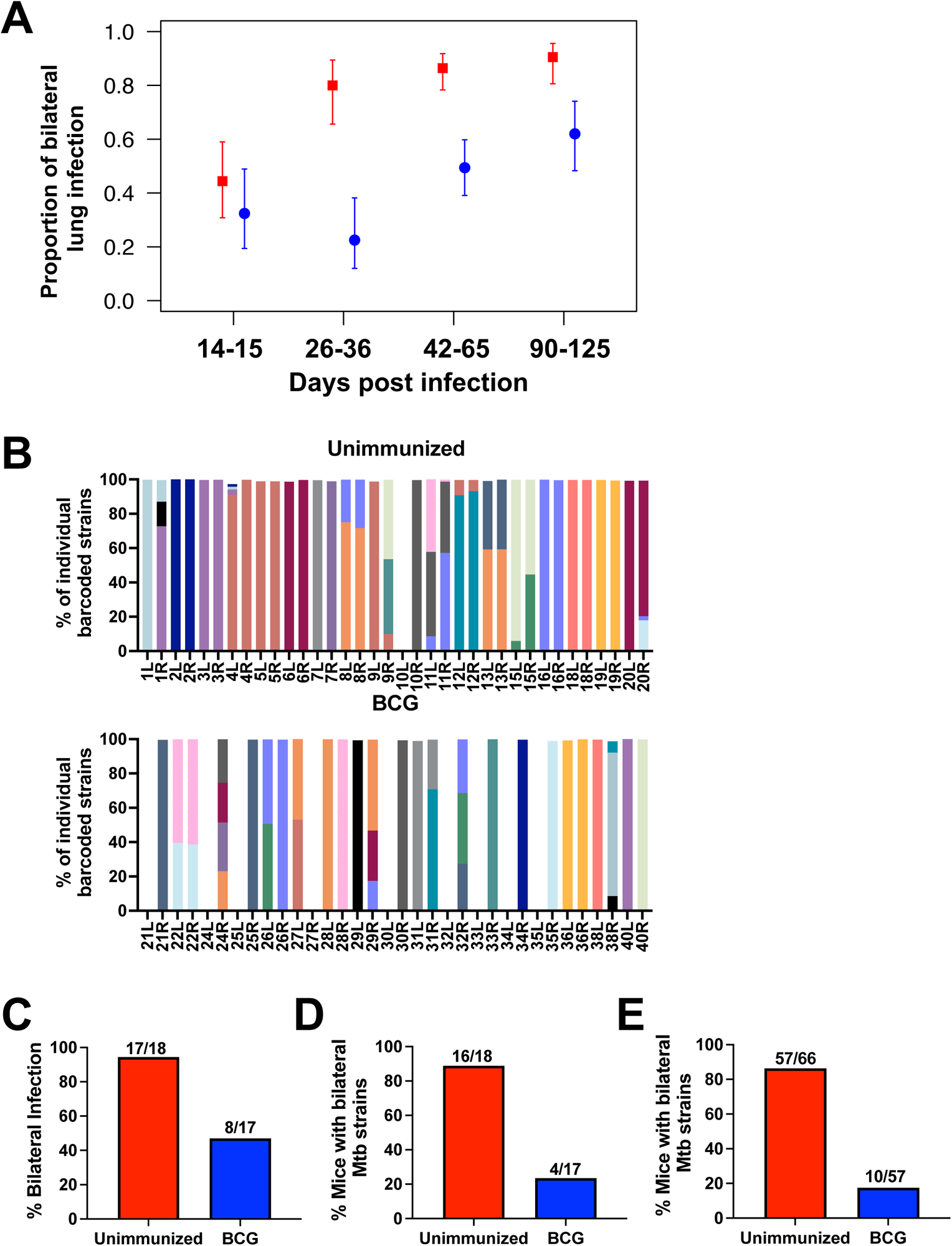
BCG-immunization prevents Mtb dissemination to the contralateral lung. **A)** Proportion of mice with bilateral lung infection (CFU in left and right lung) from a compilation of 31 experiments separated by timepoint post-infection of ULD-infected B6 mice with or without BCG immunization. Error bars are 95% confidence intervals in a mixed effects logistic regression model, with experiment as a grouping variable. All timepoints except days 14-15 post-infection have a p ≤ 0.001. **B)** A single ULD experiment using bar-coded Mtb strains is shown. On day 65 post-infection, right and left lung homogenates were plated onto 7H10 plates and Mtb colonies from infected lungs were scraped to make genomic DNA. DNA was sequenced, and the identity of each bar-coded Mtb strain is graphed for each lung separately. The percentage of bilateral infection for unimmunized and BCG-immunized mice from this experiment was calculated by the proportion of mice with CFUs in both lungs **(C)** or the proportion of mice with at least one common Mtb strain in both lungs **(D)**. **(E)** The percentage of mice with bilateral Mtb strains was compiled from 5 independent experiments, excluding day 14 post-infection. Vaccine efficacy for preventing dissemination to the contralateral lung was calculated as 1-(% BCG mice with bilateral infection)/(% Unimmunized mice with bilateral infection).

### BCG immunization can prevent detectable infection

Finally, we compared the proportion of unimmunized and BCG-immunized mice that presented with undetectable pulmonary infection. Pooling the data from all experiments assessed at D14-15 after Mtb challenge (n=6), we observed no difference in the proportion of mice with undetectable bacterial burdens in the BCG-immunized vs. unimmunized groups (**Figure 5A**). However, at all later timepoints <u>through day 125 post-infection</u>, we observed more mice with undetectable bacterial burdens in BCG-immunized mice than in unimmunized mice. Excluding the six experiments assessed at D14-15 when no difference was seen, we plotted the remaining 25 experiments at timepoints from D26-125 as the proportion of mice with undetectable bacterial burdens in the unimmunized vs. BCG-immunized group (**Figure 5B**). Although only three experiments (15-20 mice/group/experiment) reached statistical significance on their own (all in the BCG-immunized group), most experiments (18 of 25) had a higher proportion of mice with undetectable bacterial burdens in the BCG-immunized group. When the compiled data from all 25 of these experiments were assessed (**Figure 5B**, black filled circle), the difference in the proportion of mice with undetectable bacterial burdens in the BCG-immunized compared to the unimmunized group was relatively modest (13% efficacy in preventing detectable infection), but highly statistically significant (p = 0.001).

**Figure 5.**
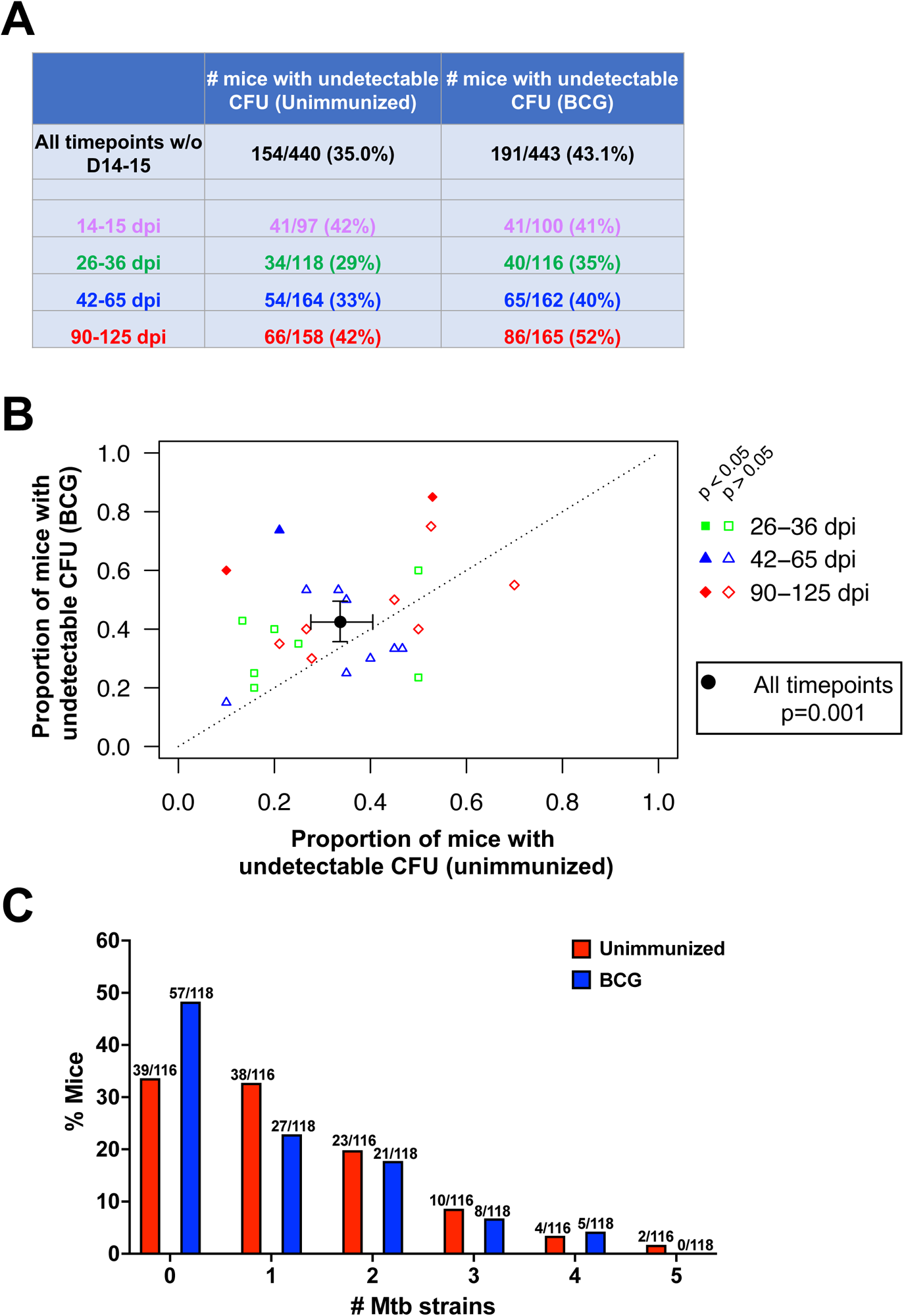
BCG immunization prevents detectable infection in some mice. **A)** Table of all mice, showing the percentage with 0 CFU in the unimmunized vs BCG-immunized groups separated by timepoint post-infection. **B)** The proportion of unimmunized mice with 0 CFU (x-axis) vs the proportion of BCG-immunized mice with 0 CFU (y-axis) for each experiment from a compilation of 25 experiments (days 26-125) separated by timepoint post-ULD infection. Each colored symbol is an independent experiment, and the larger black circle is the compilation of all the data, which was statistically significant (mixed effects logistic regression p = 0.001). Post-hoc analyses indicated that, if each infection cohort had been analyzed separately, the 3 filled in symbols would have attained statistical significance if they had been analyzed using the same regression model (p<0.05), see supplemental Figure 1. **C)** In ULD-Mtb experiments in which bar-coded strains were used for infection (n=9 for unimmunized, n=6 for BCG-immunized), the mice are graphed according to the number of unique Mtb strains recovered from each mouse’s lungs.

We also examined the number of founding strains in each mouse for all ULD experiments using the pool of bar-coded strains (9 experiments with unimmunized mice and 6 with BCG-immunized) (**Figure 5C**). As expected from the Poisson distribution, most unimmunized mice were infected with only one strain, and fewer mice were infected with two or more strains. There were similar proportions of unimmunized and BCG-immunized mice infected with two or more Mtb strains. However, fewer BCG-immunized mice were infected with one founding Mtb strain (36.9% of unimmunized mice versus 22.9% of BCG-immunized mice, p = 0.011), whereas more BCG-immunized mice had zero Mtb strains (33.0% of unimmunized mice versus 48.3% of BCG-immunized mice, p = 0.008). This suggests that infection attributable to a single founding Mtb strain may be more readily prevented by BCG-mediated immunity than infection due to two or more strains, but further investigation is needed to test this possibility more rigorously.

The demonstration that vaccine-induced immunity can prevent detectable Mtb infection in ULD-infected mice opens the possibility of assessing a new and important immune parameter for TB vaccine testing that was not known to be achievable in mice. However, BCG had a very low capacity to mediate this type of protection (13% efficacy, p = 0.001); large numbers of mice were required to reach statistical significance, which would not be practical for routine testing of TB vaccine candidates. Because the overarching goal is to identify vaccines that are more efficacious than BCG, we performed a power calculation to determine what level of vaccine efficacy would be required to feasibly measure prevention of infection with a reasonable number of mice (**Table 1)**. This analysis showed that a vaccine with 50-60% efficacy could be assessed with a sample size of 28-55 mice per group, which would be achievable by pooling results from 2-3 experiments. Such a pipeline could be feasible, as Vidal et al. recently reported that a novel live attenuated Mtb vaccine was dramatically more effective than BCG in the ULD challenge model, and prevention of detectable infection could be measured in a statistically significant manner with only 18 mice per group^25^. Taken together with our findings, these results suggest that the ULD challenge model provides a larger window to measure differences between TB vaccine candidates and to measure parameters of protection, including inhibition of dissemination and prevention of detectable infection, that cannot be assessed in currently used mouse models.

**Table 1.** Group sizes needed to assess vaccine-mediated prevention of detectable infection. Minimum sample size required per group for specified power to detect a given vaccine efficacy (prevalence in unimmunized mice assumed to be 61.6%).

|  |  | Min. sample size per group |  |
| --- | --- | --- | --- |
| Prevalence | Vaccine efficacy | 80% Power | 90% Power |
| 61.6% | 20% | 259 | 342 |
| 61.6% | 30% | 112 | 155 |
| 61.6% | 40% | 66 | 84 |
| 61.6% | 50% | 40 | 55 |
| 61.6% | 60% | 28 | 37 |
| 61.6% | 70% | 21 | 25 |
| 61.6% | 80% | 16 | 18 |
| 61.6% | 90% | 12 | 16 |

## DISCUSSION

Despite widespread vaccination with BCG, TB remains a major cause of global morbidity and mortality^14^. Although surpassed for a couple years by SARS-CoV-2 as the leading cause of death due to a single infectious organism, recent mortality estimates for each infection suggest that TB is again the number one cause of infectious death^14^. New vaccines that provide better efficacy than BCG are needed. To triage the growing number of TB vaccine candidates and move those with the most promise into clinical trials, there is an urgent need to develop small animal models that reliably assess parameters of immunity with relevance to human protection. There is growing concern that the current mouse model, in which mice are infected with 50-100 Mtb CFU by aerosolization, is not up to this task^6, 7^. This model provides too small a window to discern differences between vaccine candidates; most confer a transient reduction in the lung bacterial burden by about one log if measured between 4-6 weeks post-infection. Durable reductions in lung bacterial burdens and other clinically relevant parameters of immunity, including the ability to curb Mtb dissemination or prevent detectable infection, are difficult or impossible to assess. Indeed, the inability of the current model to predict how well vaccine candidates will perform in human efficacy trials is becoming increasingly apparent^6, 7^. In this study, we demonstrate that the ULD murine model provides a promising new challenge model in which three distinct parameters of protection can be assessed: 1) reductions in lung bacterial burdens <u>that are more durable than after conventional dose challenge</u>, 2) inhibition of Mtb dissemination, and 3) prevention of detectable infection.

Of these, reduction in bacterial burdens is the only parameter of protection that can readily be assessed in current mouse models. In contrast to transient protection that dissipates within 3-4 months in the current model^8, 9^, we have shown durable lung bacterial burden reductions for at least four months in the ULD model. Importantly, the types of immune responses that mediate reductions in lung bacterial burdens in mice infected with high doses (>250 CFUs) are sometimes different than those that do so at conventional doses (50–100 CFU)^15, 16^. Although this has not yet been rigorously assessed in the ULD model, it is reasonable to hypothesize that the optimal immune responses that are optimal for reducing lung bacterial burdens after a physiologic dose challenge of 1-3 CFU may also be different than those required for a 50-100 CFU challenge. Thus, vaccine candidates may vary in their capacity to control bacterial burdens to a physiologically relevant challenge dose compared to a challenge dose that is artificially high, a hypothesis that needs further investigation.

The second parameter of immunity that can be measured in the ULD model is the ability to prevent dissemination. Because most ULD-infected mice are infected by a single founding bacillus^11^, infection is usually initiated at a single site in one lung. Thus, most bilateral lung infection is the result of Mtb dissemination from the initially infected lung to the contralateral lung and CFU determinations of each individual lung can provide an estimation of a vaccine’s ability to block dissemination, which we have termed containment. This approach underestimates containment, however, because some mice may have Mtb in each lung not because of dissemination, but due to independent infection events of each lung by two or more distinct aerosolized strains. By infecting mice with a pool of bar-coded strains, we have shown that we can distinguish between disseminated infection and separate infections with different strains by sequencing the Mtb bar-codes in each lung. Using this approach, we have shown that BCG immunization can prevent Mtb dissemination to the contralateral lung in ∼80% of ULD-infected mice. These results in the murine ULD model parallel results in BCG vaccinated humans showing that BCG vaccination is most effective at preventing disseminated forms of TB^17^.

The third parameter of immunity that can be assessed in the ULD model is the prevention of detectable infection. There are several lines of evidence that the human immune system can prevent or eradicate Mtb infection. Some individuals with high exposure to index cases with active TB disease, including household contacts, fail to develop an Mtb-specific IFNγ-producing T cell response, suggesting that long-term Mtb infection is sometimes not established despite intense exposure^18^. Furthermore, many individuals that do develop a T cell IFNγ response against Mtb may eventually eradicate infection, as the incidence of active TB is quite low amongst IGRA+ individuals (usually less than 5%) even when their immune systems are potently immunosuppressed or ablated^19^. Recently there have been several high-profile TB vaccine studies showing that some vaccines can prevent detectable infection in non-human primates^20–22^. This had not been previously shown in mice, and it has been postulated that mice lack the fundamental immune effectors needed to prevent sustained Mtb infection^10^. In this study we showed no difference in the proportion of BCG-immunized mice with detectable infection compared to controls at d14-15 after Mtb challenge, suggesting that vaccination did not block the initial Mtb infection. At all later timepoints, however, BCG vaccinated animals had a modest, but highly statistically significant increase in the proportion of mice with undetectable infection compared to unimmunized controls (overall 13% efficacy, p=0.001). This suggests that a small proportion of the vaccinated mice that may have been initially infected were able to clear Mtb to undetectable levels. Even though BCG can do this only modestly, these results suggest that vaccine-mediated immunity can prevent sustained infection in mice exposed to a physiologic Mtb dose, challenging the longstanding belief stemming from experiments with an artificially high Mtb exposure dose, that mice are unable to eradicate Mtb infection.

One limitation of the ULD model for vaccine testing is the number of animals that are required in each group. This is exacerbated by the fact that not all animals are initially infected in the model, and currently it is not possible to discern animals that were never infected from those that were initially infected, but subsequently eradicated Mtb. We have attempted to develop both immunologic and molecular assays to distinguish between these possible outcomes, but this has proven difficult to achieve. We initially assessed Mtb-specific CD4 T cell responses against an Mtb antigen (ESAT-6) that is not present in BCG. We identified a couple unvaccinated mice that were exposed to aerosolized ULD Mtb and had measurable Mtb ESAT-6-specific CD4 T cell responses despite having no detectable lung bacterial burdens. These results suggested that even a few unvaccinated ULD-infected mice may clear Mtb to undetectable levels, but mice exhibiting this phenotype were rare. However, this approach was not successful in BCG-immunized mice because BCG immunization suppressed the development of Mtb ESAT-6-specific T cells to undetectable or almost undetectable levels even in ULD Mtb-challenged mice that were demonstrably infected, providing minimal window to discern differences between uninfected and infected mice. We also attempted to amplify Mtb DNA from lung homogenate using previously published Mtb-specific PCR primers^23^. Unfortunately, the sensitivity of this assay was not sufficient to reliably detect below 1,000 viable bacteria, and we were unable to obtain a signal from mice with undetectable bacterial burdens. Despite our inability to differentiate mice that were never infected from those who cleared infection, we were able to build strong statistical evidence for prevention of detectable infection by assessing large numbers of mice.

Our results are consistent with findings that BCG can sometimes provide long-term protection against human TB, and are consistent with studies showing BCG-mediated protection in individuals without prior Mtb exposure (tuberculin skin test-negative or IGRA-negative individuals) or in low Mtb transmission settings^1–4^. The low overall observed efficacy (∼13%) in preventing detectable infection also reflects the suboptimal nature of BCG-mediated immunity. Reliably assessing this parameter of protection for a vaccine with such low efficacy would require hundreds of mice per group, as in this study, which would not be feasible for routine pre-clinical evaluation of TB vaccine candidates. However, the goal is to identify promising vaccine candidates that are significantly more efficacious than BCG to move into human trials. Our power analysis showed that a vaccine with 50% efficacy could be readily assessed by repeating studies with 15-20 mice per group 2-3 times and compiling the results. We believe this is feasible and would be worthwhile if further studies show that results obtained in the ULD model are superior to those obtained in the conventional murine model for distinguishing vaccine efficacies in clinical meaningful ways. We are encouraged that a recent study assessing a novel TB vaccine candidate (ΔLprG, a live-attenuated Mtb vaccine) in the ULD mouse model showed that ΔLprG was dramatically better than BCG at preventing detectable infection and achieved statistical significance with only 18 mice per group^25^. In this same study, ΔLprG was only slightly better than BCG in reducing lung bacterial burdens in mice challenged with 100 CFU, suggesting that the ULD challenge model provides a larger window to discriminate differences between vaccines.

Overall, the ULD challenge model holds promise as a new and improved platform for evaluating TB vaccine candidates. The model can assess distinct parameters of vaccine-mediated immunity that cannot be assessed in the current mouse model and has potential to improve discrimination between the protective capacities of different vaccines. Each of the three parameters of immunity that can be assessed in the ULD model may be relevant to different clinical TB outcomes. For example, the ability of vaccines to durably reduce lung bacterial burdens and prevent dissemination may reflect their capacity to prevent different aspects of TB disease, whereas the ability to prevent detectable infection may reflect prevention of sustained infection. Because each of these parameters are likely mediated by different aspects of immunity, it is possible that different vaccines will differ in the relative capacity to control Mtb burdens, inhibit dissemination, and prevent detectable infection. Future studies are needed to assess a variety of TB vaccine candidates in the ULD model, and whenever possible, determine whether the results correlate with clinical outcomes in human vaccine trials.

## ACKNOWLEDGMENTS

The authors would like to thank Bridget Alexander, Lindsay Engels, Samuel Schrader, Kaitlin Durga and the Seattle Children’s OAC staff for technical support. This work was supported by NIH contract 75N93019C00070 (K.B.U), the Bill and Melinda Gates Foundation, INV-026296 (K.B.U) and NIH grant U19AI135976 (K.B.U).

## METHODS

### Mice

C57BL/6J mice were purchased from Jackson Laboratories (Bar Harbor, ME). Female mice between the ages of 9-12 weeks were used. All animals were housed and maintained in specific-pathogen-free conditions at Seattle Children’s Research Institute (SCRI). All animal studies were performed in compliance with the SCRI Animal Care and Use Committee.

### BCG Immunizations

BCG-Pasteur was cultured in Middlebrook 7H9 with OADC supplement and 0.05% Tween-80 at 37°C with constant agitation for five days. BCG was back diluted in 7H9 for two days and grown to an OD of 0.2-0.5. Bacteria was diluted in PBS and mice were injected subcutaneously with 200µl of 10^6^ CFU. After immunization, mice were rested for 8 weeks prior to Mtb infection.

### ULD Mtb Aerosol Infections

H37Rv or bar-coded H37Rv Mtb were used for infections^11^. Mtb stocks were grown in Middlebrook 7H9 with OADC supplement and 0.05% Tween-80 at 37°C with constant agitation to an OD = 1. Cultures were filtered through a 5µm filter to remove clumps and aliquots were frozen at -80°C. Frozen filtered stocks were thawed and titered side by side with stocks used for conventional dose infection to determine how to dilute the ULD stocks with the goal of leaving 37% of mice uninfected. Mice were infected using a Glas-Col aerosol infection chamber.

### CFU Plating

Mouse organs (right lung, left lung, or spleen) were homogenized separately in M tubes containing 1mL PBS+0.05% Tween-80 (PBS-T) using a Miltenyi GentleMACS machine (Miltenyi). Homogenates were then diluted in PBS-T and plated onto 7H10 plates. For ULD infections, undiluted homogenate was also plated between two 7H10 plates. Plates were incubated at 37°C for at least 21 days before quantification of CFU.

### Genomic DNA Extraction

Bacterial colonies grown from infected left lungs or right lungs were scraped into resuspension buffer (25mM Tris-HCl pH 7.9, 10mM EDTA, 50mM glucose, water) plus 10mg/mL lysozyme and were incubated at 37°C overnight. Samples were resuspended in 10% sodium dodecyl sulfate and 10mg/mL Proteinase K and were heated at 55°C for 30 minutes. Samples were then resuspended in 5M NaCl followed by Cetrimide saline solution and heated at 65°C for 10 minutes. Genomic DNA was extracted twice with 24:1 chloroform:isoamyl alcohol. DNA was precipitated with 0.7x volume of isopropanol and washed with 70% ethanol. Finally, DNA was eluted with DEPC water.

### Barcoded Sequencing

Mice were infected with a pool of 50 bar-coded strains. Sequencing of bacterial bar-codes has been previously described^11, 24^. Briefly, genomic DNA was pre-amplified with pooled barcoded primers before libraries were prepared with NEBNext Ultra DNA Library Prep Kit for Illumina (New England Biolabs) using the AMPure XP reagent (AgenCourt Bioscience) for size selection and cleanup. The NEBNext Multiplex Oligos for Illumina (New England Biosciences) were used to barcode DNA libraries and enabled multiplexing of 96 libraries per sequencing run. Samples were sequenced using the NextSeq 500 Mid Output v2 kit (Illumina) at the University of Washington Northwest Genomics Center. Read alignment was carried out using a custom processing pipeline that has been previously described^24^.

### Statistics

All statistical analysis was done in R v.4.2.0 with packages Exact (v3.1) and lme4 (v1.1-30). When comparing values between two groups in a single experiment, we used Barnard’s exact test for differences in proportions and simple linear regression on log-transformed CFU values for differences in bacterial burden (<u>excluding those with 0 CFU)</u>. For analyses compiling more than one experiment, we used mixed effects logistic regression with experiment as the grouping variable for differences in proportions, and mixed effects linear regression on log-transformed CFU values for differences in bacterial burden. In all analyses, mice were considered protected when the CFU was undetectable in both lungs, and analyses of dissemination and overall bacterial burden were performed conditional on absence of protection.

## Notes

### Competing Interest Statement

The authors have declared no competing interest.

### Summary of Updates

This version had been revised to address reviewer's comments.

